# Beyond Equilibrium Ensembles: Time Rescaling in Coarse-Grained Simulations across Single-Molecule and Condensate Regimes

**DOI:** 10.64898/2026.09.01.748708

**Authors:** Soundhararajan Gopi, Hanling Qin, Robert B. Best, Benjamin Schuler

## Abstract

Residue-level coarse-grained simulations provide a powerful route for modeling biomolecular condensates over length and time scales that are difficult to access with atomistic molecular dynamics. Coarse-grained models have been shown to reproduce many aspects of equilibrium phase behavior. However, it remains unclear to what extent such models can reproduce the relative timescales of molecular dynamics. Here, we examine this question for complex coacervates with markedly different dynamics, formed by the highly acidic intrinsically disordered protein prothymosin α with four cationic partners: linker histone H1, protamine, polylysine, and polyarginine. Coexistence simulations using a residue-level coarse-grained model reproduce key equilibrium observables from experiments, including dense-phase concentrations, ionic-strength-dependent phase behavior, and chain dimensions in the dense and dilute phases. Dynamics are accelerated in these simulations, but a composition-specific time-rescaling factor captures the ionic-strength dependence of chain reconfiguration times within a given complex coacervate. In contrast, time rescaling is not transferable between dense and dilute phases or across condensate compositions and can depend on the chosen observable. These results show that agreement with measured equilibrium observables does not imply a universally transferable timescale for conformational dynamics in residue-level coarse-grained simulations. However, we find that the required time rescaling strongly correlates with the interaction energy of the protein chains, suggesting that the missing frictional effects arise from protein-protein interactions rather than solely from protein-solvent interactions, reminiscent of internal friction. Our findings highlight the need to combine thermodynamic validation with kinetic calibration when interpreting chain relaxation, molecular diffusion, and material properties from residue-level coarse-grained simulations of biomolecular condensates.

## **I.** INTRODUCTION

Biomolecular condensates are mesoscale assemblies of cellular importance whose properties emerge from multivalent interactions often involving intrinsically disordered proteins (IDPs)^1,2^. Molecular simulations are powerful approaches for understanding the molecular-scale conformations and dynamics essential to elucidating the mechanisms underlying the biological functions of condensates. Despite the crowded molecular environment, IDPs often retain rapid conformational dynamics within the condensate phase that give rise to distinctive material properties, molecular organization, and translational diffusion in the dense phase^3,4^. The conformational dynamics of IDPs in condensates can occur on timescales of hundreds of nanoseconds to a few microseconds, making them accessible to atomistic molecular dynamics simulations^3–9^. When experimentally validated, such simulations provide the spatial and temporal resolution needed to connect atomic-level molecular properties, at nanometer length scales and nanosecond timescales, to the mesoscale behavior of biomolecular condensates, which emerges at micrometer length scales and up to seconds timescales. However, probing the equilibrium between phases still remains beyond the reach of such simulations.

Over the past decades, implicit-solvent coarse-grained (CG) simulations, particularly residue-level CG models^10–16^, have started to provide an important bridge between atomistic simulations and polymer theory applied to biomolecular condensates. By reducing the number of degrees of freedom while retaining key molecular features, these models enable simulations of larger biomolecular assemblies over longer timescales than in all-atom (AA) simulations. Residue-level CG models have been particularly successful in describing the conformational ensembles of IDPs and their responses to changes in solution conditions, residue-level perturbations, and binding partners^11,17–21^. In these models, solvent-averaged non-bonded interactions are commonly described using a screened Coulomb potential for electrostatic interactions, in the simplest case with integer charges assigned to charged amino acids, together with residue-specific non-electrostatic interactions represented by potentials such as the Ashbaugh–Hatch^10,22^ or Wang-Frenkel^23,24^ forms. This combination provides an effective approximation of residue-level interaction energies often sufficient to capture sequence-specific effects in IDPs, and it has been shown to reproduce key equilibrium properties of phase-separating systems, including critical temperatures, phase diagrams, and the dense- and dilute-phase behavior of self-associating proteins and charged coacervates^13,18,25^.

However, the approximations that make CG models efficient also introduce limitations. In residue-level CG models, amino acids are usually represented as spherical beads with isotropic interaction fields that depend only on inter-residue distance^26–28^. As a result, a given residue can interact with more than one neighboring residue with comparable interaction energy at the same time. This differs from all-atom simulations, where side-chain-specific interactions are often anisotropic, orientation-dependent, and more strongly constrained by local geometry. The isotropic approximation, therefore, does not explicitly resolve fast molecular fluctuations such as side-chain rotations, reorientations, and short-lived contact rearrangements. Instead, these degrees of freedom are represented by an effective potential of mean force over unresolved side-chain and solvent degrees of freedom. In this sense, the isotropic interactions between spherical CG residues approximate the statistical average over side-chain configurations that interconvert on timescales much faster than the global chain-reconfiguration time^26,29^. As a result of such coarse-graining, CG simulations generally do not reproduce absolute molecular timescales and require an empirical rescaling relative to experiments or atomistic explicit-solvent simulations^30–35^.

All-atom simulations of complex coacervates formed from highly charged IDPs indicate that side-chain contact lifetimes can be orders of magnitude shorter than chain reconfiguration times^3,4^. This raises a central question: how does coarse-graining fast, anisotropic side-chain fluctuations into effective isotropic interactions affect the interpretation of equilibrium and dynamic properties derived from CG simulations? Can a CG model that reproduces equilibrium phase behavior also reproduce molecular dynamics with appropriate time rescaling, or does coarse-graining renormalize time in a system-specific way that depends on the environment in the dense versus dilute phase and on sequence composition? This latter issue arises when microscopic states are lumped into macrostates, because the resulting reduced dynamics need not remain Markovian^36^. In general, projecting onto a reduced set of coarse variables yields dynamics with fewer degrees of freedom, in which the unresolved degrees of freedom appear as a memory kernel and a fluctuating noise term^37–39^. Even if the coarse dynamics is well-approximated as being Markovian, the effective friction from the missing degrees of freedom is unlikely to be captured by the single friction coefficient commonly used in Langevin dynamics simulations. Rather than asking only whether a CG model reproduces phase diagrams, we ask whether the same model also preserves the timescales that govern processes such as chain reconfiguration and translational diffusion. This distinction is essential because equilibrium agreement can arise from effective thermodynamic averaging over unresolved atomistic degrees of freedom, whereas dynamics depend on how those averaged degrees of freedom renormalize friction, contact lifetimes, and memory effects^28,34,40–43^.

In this work, we systematically dissect the consequences of coarse-graining for charged biomolecular condensates. We simulate four charged complex coacervate systems composed of the negatively charged IDP prothymosin α (ProTα; z=-44) paired with four different cationic partners: the lysine-rich linker histone H1 (z=+54), its arginine-rich functional analog protamine (z=+22), and two synthetic homopolymers composed of lysine or arginine residues, K50 (z=+50) and R50 (z=+50), respectively [Table S1]. This set of systems has recently been investigated in detail experimentally and, for subsets of these systems, directly compared with all-atom explicit-solvent simulations^3,4^, thereby allowing us to compare natural and synthetic polycationic partners and enabling complementary comparisons that isolate some effects of charge patterning, residue chemistry, and interaction specificity. We test whether a residue-level CG model, parameterized using equilibrium observables, also preserves a transferable timescale for conformational dynamics across condensates with different residue chemistries. Our results show that residue-level CG models can capture the equilibrium properties of the dilute and dense phases, such as their protein concentrations and IDP dimensions, but they rescale biomolecular dynamics in a nontrivial, system-dependent manner. We show that CG time rescaling is not transferable but becomes composition- and environment-dependent, and can be observable-dependent. Since the required time rescaling correlates with the coarse-grained chain interaction energy, the missing frictional effects are likely due to protein-protein interactions.

## **II.** MODELS AND METHODS

Proteins were modeled at a single-bead-per-amino-acid resolution, with each bead representing an amino acid centered at its Cα position. The total potential energy (*U*) was expressed as the sum of bonded, electrostatic, and non-bonded short-range interactions.

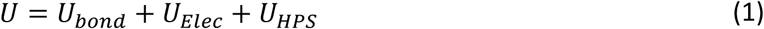

The beads are connected using a harmonic bond potential (*U_bond_*) with a force constant of 418.4 kJ nm^-2^ mol^-1^ and an equilibrium bond length of 0.38 nm. Electrostatic interactions were modeled using a screened Coulomb potential,

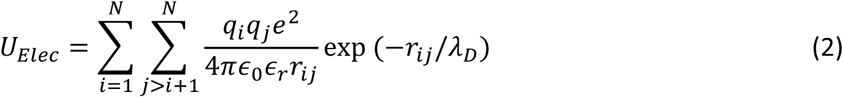

where *N* is the total number of residues, *q_i_* and *q*_j_ are the integer charges on the residues *i* and j, respectively, *e* is the elementary charge, ∈_0_ is the permittivity of free space, ∈*_r_* is the relative permittivity of water and is set to 80, *r_i_*_j_ is the distance between residues *i* and j, and *λ_D_* is the Debye screening length that depends on the ionic strength and temperature according to

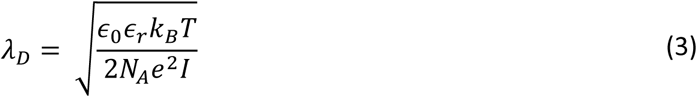

Here, *k_B_* is the Boltzmann constant, *T* is the temperature, *N_A_* is Avogadro’s constant, and *I* is the ionic strength. The non-electrostatic pairwise interactions are modeled using a short-range hydrophobicity-scale (HPS) potential that includes parameters for explicit dyes^10,11^:

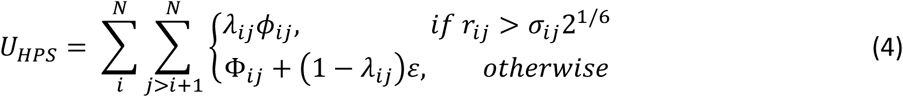

Here, *λ_i_*_j_ is the pairwise attractive interaction defined by the CG model, *σ_i_*_j_ is the excluded volume radius, *ε* is the interaction strength, and is set to 0.837 kJ/mol, and *φ_i_*_j_ is defined as

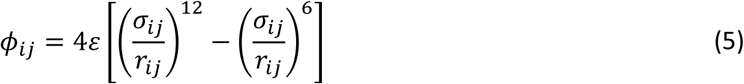

For ProTα and H1–ProTα dimer simulations, the proteins were placed at the center of a 30×30×30 nm^3^ box with periodic boundary conditions. The donor and acceptor dyes for Förster resonance energy transfer (FRET) were explicitly represented as described before^11^, and attached at the same labeling positions used experimentally^4,44^. For the slab simulations, the individual protein chains were first compacted by setting an artificially strong pairwise interaction energy, *λ*=1, for all CG beads. The polycation and ProTα chains [Table S1] were then alternately placed on a grid at the center of a 25×25×25 nm^3^ box. The number of protein chains was chosen to achieve near charge balance – H1–ProTα (88:108), protamine–ProTα (216:108), K50–ProTα (95:108), and R50–ProTα (95:108). The system was energy-minimized and simulated for 2×10^7^ steps at 300 K, 8 mM ionic strength, and 5 bar to obtain a compact initial configuration, after restoring normal residue-specific interaction parameters^11^. The production simulation was started from the resulting compact configuration of the NPT simulation. A cubic box encompassing the compact, dense initial configuration was extended by 50 nm on both sides along the x-axis, with periodic boundary conditions to enable coexistence simulations. The interactions across periodic images have been shown to perturb system dynamics^45,46^, and we ensured that the shortest simulation box dimensions were sufficiently larger than the end-to-end distance of the individual chains to reduce such artifacts^3^. The final box dimensions differed across the four systems studied to account for system-dependent coacervate densities [Fig. 1(a) and 1(d)]. Starting from the compact initial configuration, Langevin dynamics (NVT) simulations were performed with GROMACS 2019.4^47,48^ at 300 K, with a time step of 10 fs and a friction coefficient *γ* of 0.2 ps^-1^. The non-bonded interactions were truncated at a cutoff distance of 3.5 nm. The simulations were performed for 5×10^8^ MD steps; the first 2×10^8^ steps were discarded as equilibration, and coordinates were stored every 1000 steps. An additional simulation of the H1–ProTα dense phase was set up at 128 mM ionic strength and 300 K, starting from a uniformly mixed initial configuration generated from a short equilibrium simulation at 1 M ionic strength and 300 K. To determine the dependence of equilibrium and dynamical observables on the Langevin friction coefficient, additional simulations were performed at 128 mM ionic strength over the range *γ* = 0.0005, 0.008, 0.01, 0.033, 0.1, 1, 5, 10, 30 ps^-1^, in addition to the production value *γ*=0.2 ps^-1^ starting from the equilibrated initial coordinates, with all other simulation parameters unchanged, except that simulations at the highest friction coefficients (*γ* = 5, 10, 30 ps^-1^) were extended by an additional 2 μs to improve sampling of the slower dynamics.

**FIG. 1.**
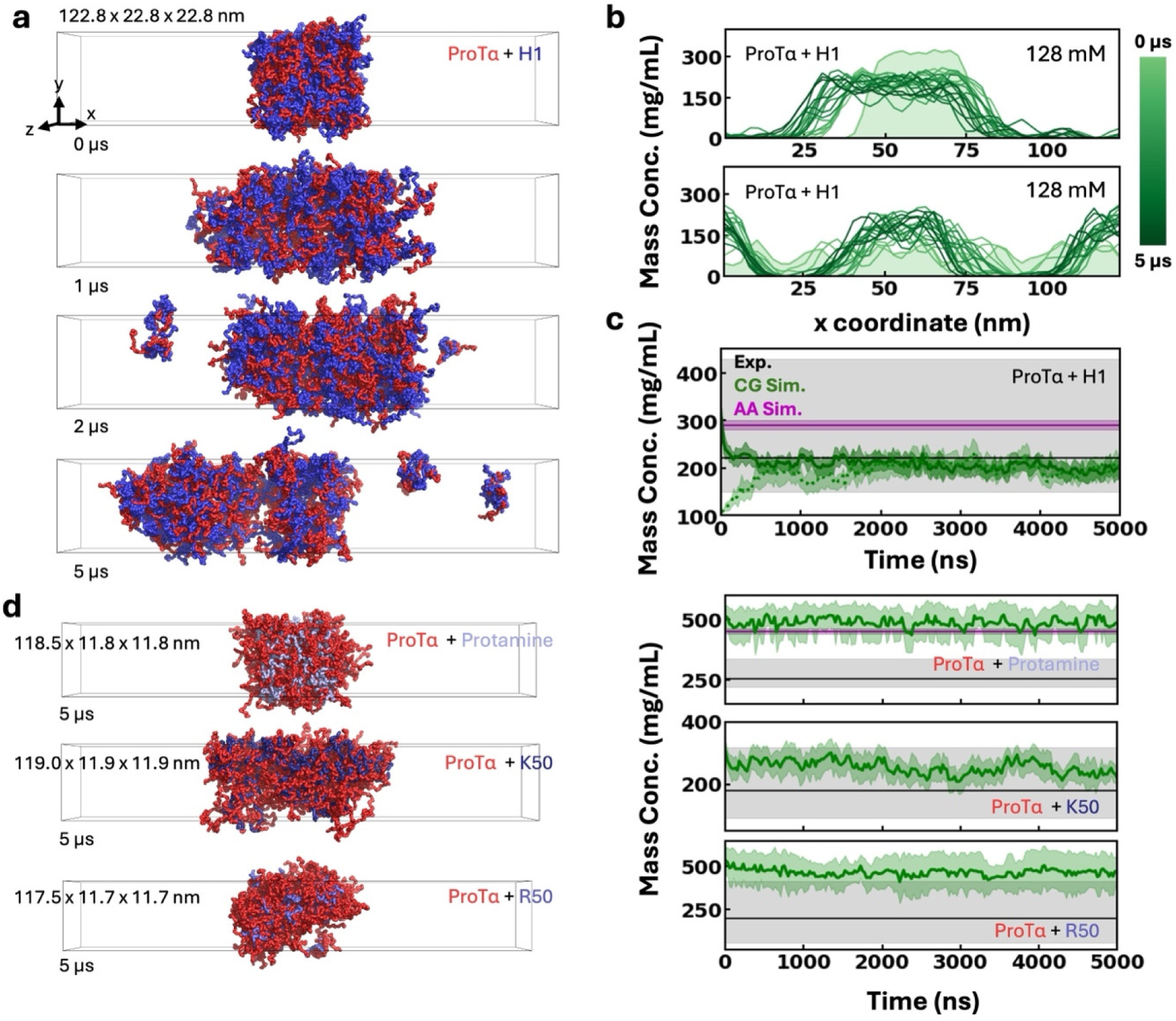
Equilibration and dense-phase protein concentrations from CG simulations. (a) Representative snapshots of the H1–ProTα condensates in slab configuration, with H1 in blue and ProTα in red, starting from the artificially dense configuration at 128 mM ionic strength. The simulation box is shown as black lines. (b) The time evolution of the density profiles along the x-coordinate (long axis of the slab), starting from the artificially condensed (top) and uniformly mixed (bottom) initial configurations. The initial density is shown as shaded green. (c) Equilibration of dense-phase protein concentration starting from the artificially dense (solid green) and uniformly mixed (dotted green) initial configurations. The concentration estimates from single-molecule experiments and all-atom MD simulations are shown in black and purple, respectively. The shaded areas indicate the associated uncertainties. (d) Representative snapshots from the CG simulations of the other three condensates (left) and the time dependence of their dense-phase protein concentrations (right). The color code is the same as in panel (c).

The density profiles along the *x*-axis were calculated over non-overlapping 2.5×10^6^ step windows to study density equilibration throughout the simulations [Fig. 1(b)]. The average dense-phase concentration was computed by first identifying the high-density region, excluding the interface, and then using the average and standard deviation of the density within this region to estimate the uncertainty for each window along the *x*-axis of the simulation box. Similarly, the dilute-phase concentration is the average concentration of the box region that never overlapped with the high-density or interface regions.

The trajectories were analyzed in Python using the MDAnalysis library^49^. The FRET efficiencies were calculated from the inter-dye distances, *r*, using the Förster equation:

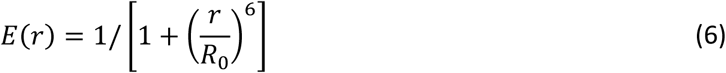

Here, *r* is the distance between the central dye beads in the ProTα and H1–ProTα dimer simulations, *R*_0_= 5.4 nm is the Förster radius of the Alexa dye pair, and *R*_0_= 6 nm for the Cy3B-CF660R dye pair used in the experiments^3,44^. In dense-phase simulations, the dyes were accounted for implicitly because the labeled-to-unlabeled protein ratio is too low to model explicitly. The dye-dye distance *r* was calculated from the Cα coordinates of the experimentally labeled positions as *r* = *d*((*N* + 9)/*N*)*^u^*, where *d* is the distance between the labeled positions, *N* is the sequence separation between the labeled positions, and the scaling exponent *υ* was set to 0.6.^3^ To assess the accuracy of this approximation, we additionally performed a representative H1–ProTα dense-phase simulation containing one explicitly labeled ProTα chain, with donor and acceptor dyes attached at residues matching the experimental setup^3^, using the same explicit-dye model as in the dilute-phase simulations^11^. For this trajectory, the dye-dye distance was calculated directly from the central dye beads. In slab simulations, *R*_0_=5.9 nm was used to account for the refractive index change in the dense phase^3^. In all cases, we assumed rapid orientational averaging of the dyes, corresponding to an orientational factor of 2/3. The chain reconfiguration time (*τ_r_*) was calculated from the simulations based on *r* from the distance correlation function

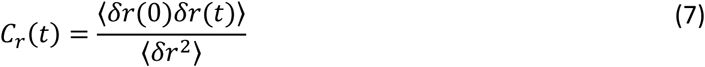

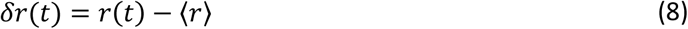

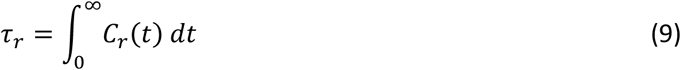

The translational diffusion coefficients (*D*) of the center of mass of individual ProTα chains were calculated using GROMACS^47,48^ after accounting for discontinuities caused by periodic boundary crossings. Because the dense phase adopts a slab geometry with the interface normal to the *x*-direction, translational motion along *x* is spatially confined by the finite thickness of the dense phase and therefore does not represent unrestricted bulk diffusion. We consequently calculated the mean-square displacement using only the two directions parallel to the slab, *y* and *z*, and obtained the lateral diffusion coefficient from the two-dimensional Einstein relation, MSD*_yz_*(*t*) = 4*D_yz_t*. The time interval between reference points used to calculate the mean square displacement (MSD) was set to 10 ns, and the MSD curves over 10-90% of the simulation time were fitted to calculate *D*. For an isotropic bulk liquid, the diffusion coefficient inferred from the two unconstrained Cartesian components is equivalent to that obtained from three-dimensional diffusion. In the slab simulations, however, exclusion of the confined *x*-direction avoids bias associated with the finite dense-phase thickness and interfaces. A comparison of diffusion estimates obtained from different coordinate projections is shown in Fig. S1.

The average transfer efficiency, (*E*⟩, reconfiguration time, *τ_r_*, and diffusion coefficient, *D*, were calculated for individual ProTα chains in the dense phase simulations, and the average and standard deviation across all chains are reported. Diffusion estimates from finite condensate trajectories can also depend on the local molecular environment, the finite simulation size, and the slab geometry. The uncertainties reported should not be interpreted as encompassing all systematic uncertainty in the absolute diffusion coefficient. For monomeric ProTα and H1–ProTα dimer simulations, uncertainties were estimated from block averages by dividing the trajectories into three non-overlapping blocks. For each friction coefficient, the time-rescaling factor (*S_t_*) was defined as the ratio of the experimental reconfiguration time to the corresponding CG simulation value. The dependence of the time rescaling factor on the friction coefficient (*γ*) was fit using an empirical shifted inverse function

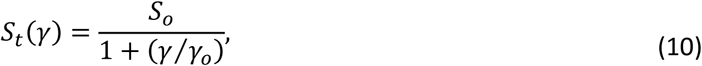

where *S_o_* is the limiting time-rescaling factor at low friction, and *γ_o_* (ps^-1^) is the characteristic crossover friction coefficient at which *S_t_* = *S_o_*⁄2. This expression approaches the correct high-friction limit *S_t_* ∝ *γ*^−1^expected for overdamped Langevin dynamics^50,51^ and models the turnover at very low friction consistent with the low-friction regime of Kramers dynamics^51^. For each ProTα chain, the interaction energy (*E*_int_) was calculated as the total non-bonded interaction energy involving that chain, including both intramolecular and intermolecular contributions, and comprising the electrostatic and HPS terms.

## **III.** RESULTS AND DISCUSSION

### **A.** Dense-phase simulations

We performed coarse-grained (CG) coexistence simulations of IDP complex coacervates using slab configurations in which one box dimension was extended relative to the others to enable simultaneous sampling of the dense and dilute phases^10,52–54^. CG slab simulations of H1–ProTα initiated from dense configurations at *γ*=0.2 ps^-1^, relaxed within ∼200 ns of CG simulation time to a protein concentration in the dense phase of approximately 220±9 mg/mL at an ionic strength of 128 mM [Fig. 1(a)-1(c)], close to the experimental estimate of 220±70 mg/mL.^3^ To assess whether the results were biased by the initial configuration, we also simulated the H1–ProTα system starting from an initial state with a protein concentration of 100 mg/mL uniform across the simulation volume. In this case, the dense phase formed spontaneously and reached the same concentration as the simulation starting from the dense phase within approximately 1 µs, supporting equilibration of the dense-phase concentration. Multiple dense-phase droplets were observed during the simulations under both initial conditions [Fig. 1(a)-1(b)]. Because their diffusion and coalescence were slow relative to the simulation timescale, these domains could persist for several microseconds without merging.

To further evaluate the performance of the CG model, including its sensitivity to charge patterning and residue-specific interactions, we performed coexistence simulations of ProTα with three other polycationic partners: protamine, K50, and R50.^4^ Linker histone H1 and protamine are naturally occurring proteins with related roles in DNA compaction^55,56^, whereas the Lys and Arg homopeptides K50 and R50 are synthetic polyelectrolytes with maximal charge density that provide simplified test systems for evaluating the treatment of local residue–residue repulsion and residue identity in CG simulations. As observed for the H1–ProTα condensate, the dense-phase concentrations of the other condensates at 128 mM ionic strength converged to values close to the corresponding experimental estimates on timescales of a few hundred nanoseconds [Fig. 1(d)]. After equilibration, fluctuations in dense-phase concentrations were small, and the uncertainties were comparable in magnitude to the experimental uncertainties. The average dense-phase concentrations of condensates containing the lysine-rich cationic partners H1 and K50 were close to the experimental values^4^. For the arginine-rich condensates, however, the dense-phase concentration was higher than experimentally observed but comparable to that obtained from all-atom molecular dynamics simulations for the protamine–ProTα condensate^4^. The remaining discrepancies between simulated and experimental dense-phase concentrations may indicate that arginine-mediated interactions are slightly too strong in both the CG and all-atom molecular dynamics models^57–60^.

### **B.** Ionic-strength-dependent phase behavior

As shown in Fig. 1, coexistence simulations of the H1–ProTα condensate spontaneously populate a dense and a dilute phase. At 128 mM ionic strength, the dilute phase contains small oligomers, predominantly H1–ProTα dimers, trimers, and tetramers, whose presence has been experimentally observed^44^. To evaluate how well the CG simulations capture ionic-strength-dependent phase behavior, we performed H1–ProTα coexistence simulations at different ionic strengths [Fig. 2(a,b)]. Consistent with the behavior expected of complex coacervates, the dense-phase concentration decreased as ionic strength increased. Equilibrated concentration profiles along the long axis of the box showed that the dense-phase concentrations converged to values close to the corresponding experimental estimates and remained stable over microseconds [Fig. 2(b)]^4^.

**FIG. 2.**
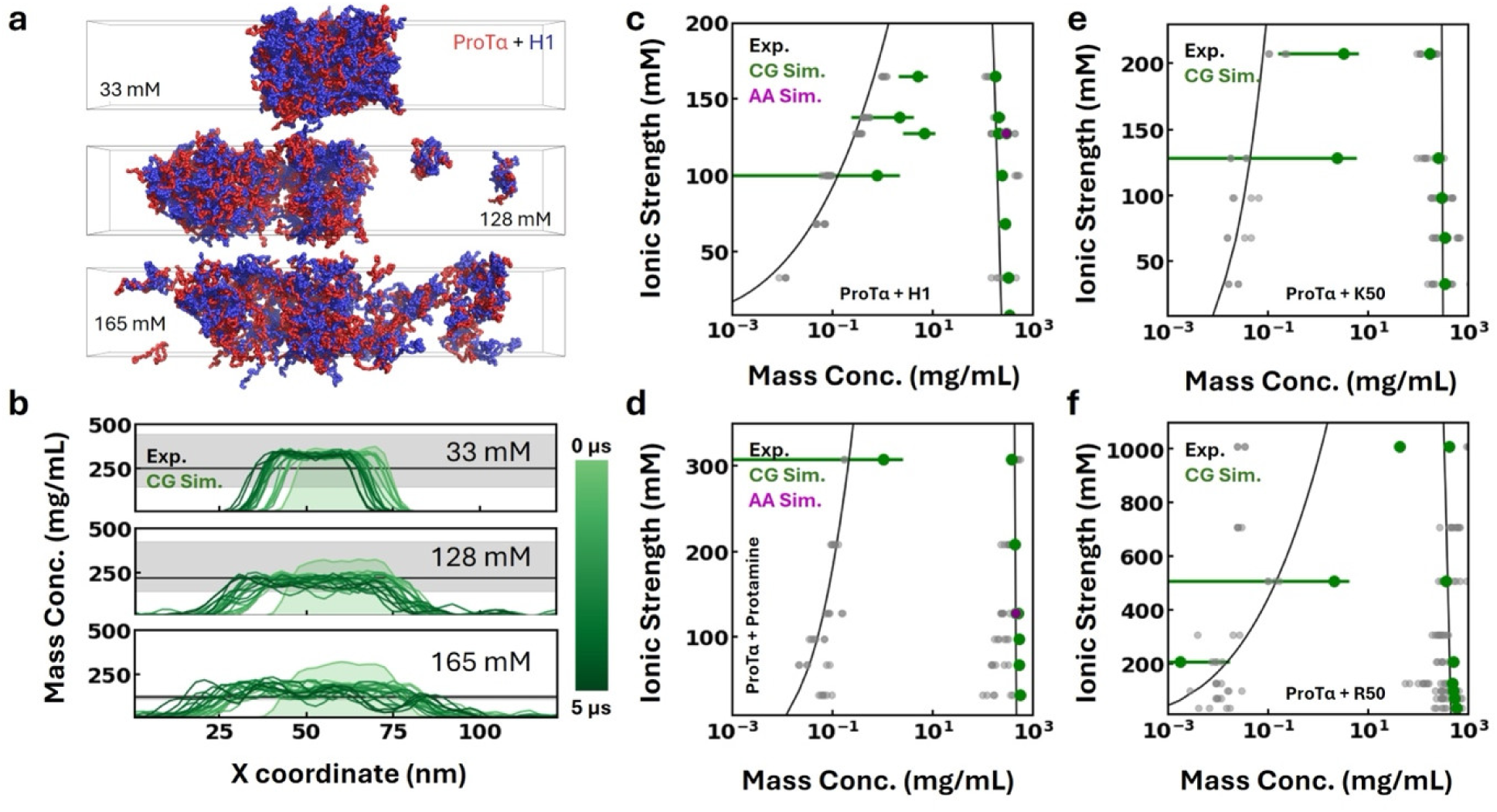
Ionic-strength-dependent phase diagrams from CG simulations. (a) Representative snapshots of the CG simulations of H1–ProTα condensate at different ionic strengths. (b) Time evolution of the density profiles along the longest box coordinate at three ionic strengths. The black line indicates the experimental dense-phase concentrations, and the shaded area represents the associated uncertainties. (c-f) Phase diagrams from the coexistence simulations as a function of ionic strength. The experimental data points are shown in gray, and the estimates from CG and AA simulations are shown in green and purple, respectively. The phenomenological fit to the experimental data, based on the Voorn-Overbeek theory^61^, is shown as black lines for reference.

To reconstruct the phase diagrams of the four systems, we performed slab coexistence simulations at ionic strengths matching the bulk salt concentrations used in the experiments^4^. Across all four systems, the simulated ionic-strength-dependent changes in dense-phase concentration were close to the experimental measurements [Fig. 2(c)–2(f)]. The accuracy is comparable to that of the Mpipi-Recharged model, despite the two models being optimized independently on distinct datasets^13,62^. Accurate estimation of dilute-phase concentrations, however, is more challenging owing to the small volume of the dilute phase and the correspondingly small number of oligomers populated in the simulations. For condensates containing lysine-rich cationic partners, oligomers in the dilute phase were observed at ionic strengths above 100 mM. For condensates containing arginine-rich cationic partners, oligomers in the dilute phase were populated only at ionic strengths above 200 mM. Dilute-phase concentrations were estimated from regions of the simulation box that did not overlap with either the dense phase or the interface, such as the regions near the box boundaries along the x-axis for the H1–ProTα condensate shown in Fig. 2(a). The same analysis volume was used across systems and ionic strength conditions to avoid bias introduced by differences in the number of sampled spatial points. The dilute-phase concentrations obtained from the CG simulations exhibited large uncertainties, but these uncertainty ranges generally encompassed the corresponding experimental dilute-phase concentrations. Because the dilute phase was mainly composed of small oligomers, particularly dimers and trimers, increasing either the box length or the number of protein chains would be expected to improve the statistical precision of dilute-phase concentration estimates^63^. Given the ability of the CG simulations to reproduce key features of the phase diagrams, we next examined the conformational ensembles of the intrinsically disordered proteins in the dense and dilute phases.

### **C.** IDP chain dimensions in the dense and dilute phase

Single-molecule FRET efficiencies provide a benchmark for evaluating whether the CG simulations accurately reproduce experimentally measured mean FRET efficiencies of IDPs both in the dilute phase^11,14,64^ and within their complex coacervates^3,4^. To evaluate whether the isotropic interactions between spherical CG beads can reproduce experimentally measured FRET efficiencies, we simulated ProTα under three conditions: as a monomer, as a dimeric complex with linker histone H1, and within a charge-balanced H1–ProTα complex coacervate composed of 108 ProTα and 88 H1 chains. For each system, we calculated the mean FRET efficiency, (*E*⟩, between residues 58 and 112 in ProTα, the positions labeled in the experiments^3,4,14,44,64^. Monomeric ProTα is highly expanded at low ionic strength, with (*E*⟩ ≈ 0.19. As ionic strength increases, ProTα becomes more compact because of reduced intrachain electrostatic repulsion, and (*E*⟩ reaches a plateau at high ionic strength, with (*E*⟩ ≈ 0.46 [Fig. 3(a)]. The simulated ionic strength-dependent trend in (*E*⟩ was close to the behavior observed experimentally^4^, as indicated by a concordance correlation coefficient of *ρ_c_* = 0.80. However, ProTα was more expanded in simulations than in experiments at low ionic strength, suggesting that local electrostatic repulsion may be overrepresented in the model under these conditions.

**FIG. 3.**
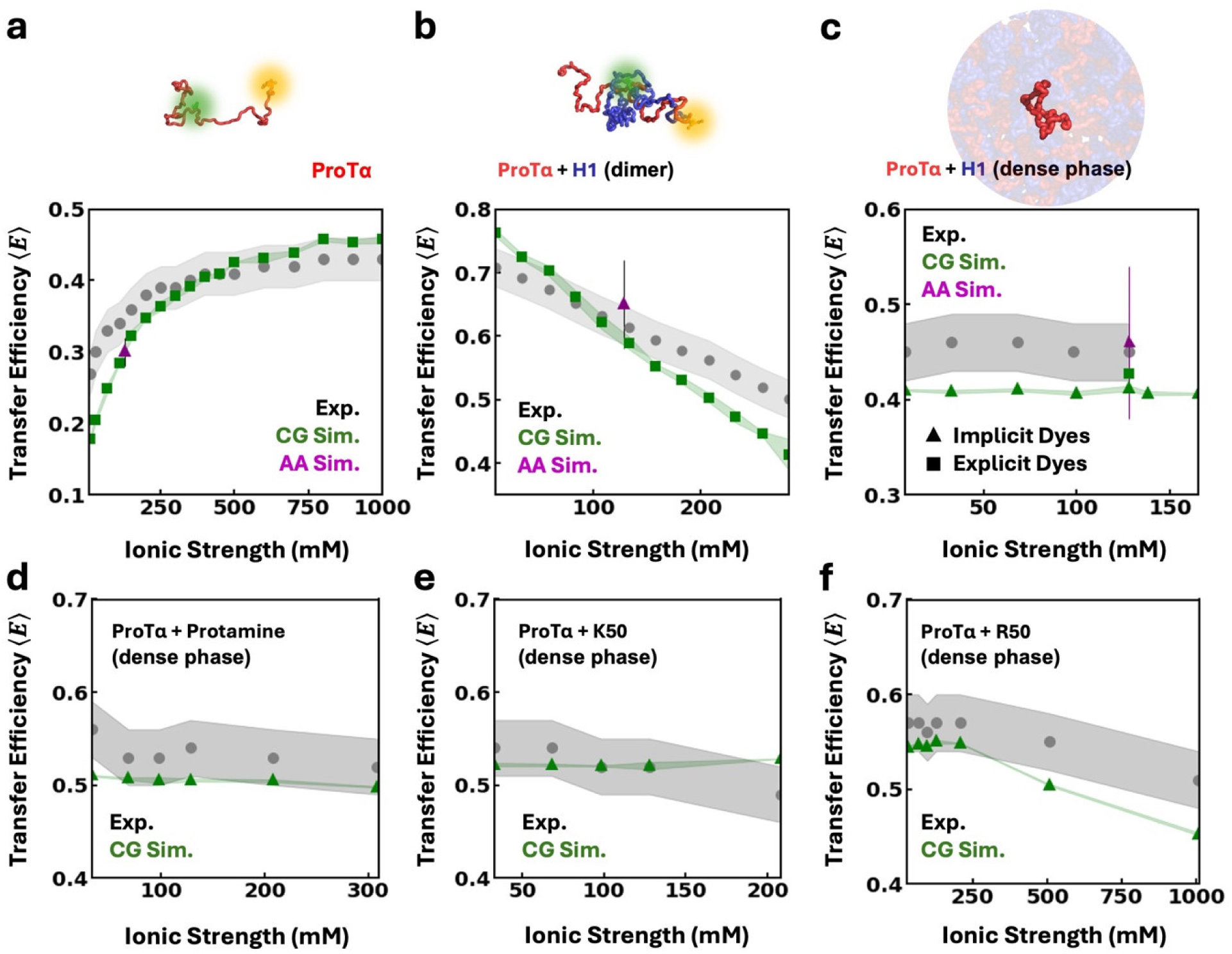
Chain dimensions in dense and dilute phases. Comparison of mean FRET efficiency, (*E*⟩, of ProTα as (a) an isolated monomer, (b) in an H1–ProTα dimer, and (c) in the dense phase of the H1–ProTα condensate. Representative snapshots of the simulated systems are shown at the top, with explicit donor (green) and acceptor (orange) dyes that match the experimental labeling positions. (d-f) Mean FRET efficiency of ProTα in the dense phases of (d) protamine–ProTα, (e) K50–ProTα, and (f) R50–ProTα condensates. Shaded bands show the uncertainties. In panels (a)-(f), triangles denote implicit-dye calculations (AA or CG), and squares denote explicit-dye simulations. Simulation uncertainties were calculated from block averages for panels (a-b) and from the distribution across ProTα chains within the condensates for panels (c-f).

In the H1–ProTα dimer simulations, the IDPs are compact at low ionic strength because of favorable interactions between oppositely charged residues on the two polyelectrolytes. With increasing ionic strength, the complex expands as electrostatic interactions are screened [Fig. 3(b)]. This ionic-strength-dependent conformational response is consistent with the experimental FRET efficiencies, with *ρ_c_* = 0.85^44^. Despite modest discrepancies at low and high ionic strengths, the simulated values were within the experimental uncertainty near the physiological ionic-strength range and comparable to estimates from all-atom simulations. In the dense phase, (*E*⟩ showed only a weak dependence on ionic strength, in agreement with the experimental observation^4^ [Fig. 3(c)–3(f)]. For the H1–ProTα condensate, a representative simulation containing one explicitly dye-labeled ProTα chain yielded a mean FRET efficiency consistent with the value obtained using the implicit-dye treatment at the same ionic strength [Fig. 3(c)], supporting the use of the implicit approximation for the dense-phase calculations. The ionic-strength dependence of ProTα chain dimensions varies substantially across molecular environments, and these distinct responses are quantitatively reproduced by the CG simulations. Isolated ProTα becomes more compact with increasing ionic strength, whereas ProTα in the H1–ProTα dimer expands over the same ionic-strength range. The chain dimensions of ProTα in the dense phase are comparatively insensitive to ionic strength over the experimentally probed range. The simulations reproduce both these changes in the direction and magnitude of the conformational response and the corresponding experimental FRET efficiencies.

Consistent with the experiment, ProTα in the R50–ProTα condensate showed a relatively stronger response to increasing ionic strength at high ionic strength, while the dense phase remained stable under these conditions. This result suggests that the CG simulations qualitatively capture the persistence of protein condensation at 1 M ionic strength, where salt-induced non-electrostatic interactions, particularly enhanced arginine hydrophobicity associated with salting-out effects^4,5,65^ that are not explicitly represented in the present force field, contribute substantially to condensate stability [Fig. 2(f) and 3(f)]. Overall, the simulations reproduce the experimental observation that ProTα adopts distinct conformational ensembles across the monomeric, dimeric, and condensate environments, including that ProTα in H1–ProTα condensates is more compact than in the monomeric state but more expanded than in the H1–ProTα dimer^3^.

Based on these comparisons alone, it is difficult to determine whether the remaining discrepancies between simulated and experimental (*E*⟩ arise primarily from the strong repulsion between neighboring residues in the CG model, from the absence of ionic-strength-dependent parameterization in the HPS interaction parameters, or from a combination of both. Additional contributions may also arise from limitations of the electrostatics treatment, neglected ion-specific and charge-regulation effects, or other approximations inherent to the coarse-grained representation. Overall, however, the HPS CG model reproduces the equilibrium conformations and protein concentrations of these four complex coacervates across the tested experimental range reasonably well, considering that the model was parameterized on an orthogonal set of disordered proteins^11^, most of which had much lower charge density. The accuracy is comparable to that obtained with a CG model explicitly optimized to match the H1–ProTα dimer FRET data^14^. We next assessed whether the simulations could capture molecular dynamics within the condensates to a similar extent.

### **D.** System-dependent time rescaling of CG dynamics and possible observable dependence

Langevin dynamics, a common approach for implicit-solvent coarse-grained simulations of biomolecules, approximates coupling to solvent and other degrees of freedom by adding a frictional dissipative force and a memoryless stochastic force to the conservative force derived from the interaction potential^38,39,48,66^.

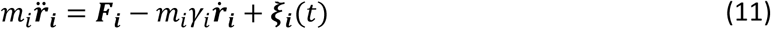

For each CG particle, the equation of motion includes the conservative force, ***F_i_*** = - Δ***U_i_***, where *U* is the total interaction potential defined in Eq. 1, a friction term (*m_i_γ_i_*) proportional to bead velocity (**ṙ***_i_*), and a random force (***ξ****_i_*) that represents thermal fluctuations. The latter two forces approximate the degrees of freedom that are integrated out during coarse-graining. The random force is constrained by the fluctuation–dissipation theorem^67^ and therefore depends on the simulation temperature, particle mass, and friction coefficient. The particle mass (*m_i_*) is defined by the CG model, whereas the friction coefficient (*γ_i_*) is chosen either to approximate solvent friction^40,68^, to provide weak thermostat coupling^10,19^, or to calibrate the dynamics to target relaxation properties^40^.

For residue-level CG models in water at 300 K, a physically motivated estimate based on Stokes friction yields a friction coefficient of *γ_i_* ≈ 30 ps^−1^, assuming a uniform residue mass of 110 Da and a hydrodynamic radius of 0.38 nm for the CG beads^69^. However, many CG simulations use a much smaller value, such as *γ_i_* ≈ 0.2 ps^−1^, to accelerate sampling and improve sampling efficiency^70^. In the overdamped limit, relaxation times are expected to increase approximately in proportion to the friction coefficient. If changing *γ* only altered this overall dynamical timescale, the corresponding experimental time-rescaling factors would decrease approximately as *γ*^−1^, but relative differences between molecular environments or compositions would remain unchanged. Because such weak-friction simulations do not directly reproduce solvent viscosity, the resulting absolute timescales must be empirically rescaled against atomistic simulations or experimental measurements^33,34^. In addition, the effect of missing protein degrees of freedom must be accounted for in the effective friction. We thus asked three related questions: First, is the time mapping transferable between molecular environments and condensates with different sequence compositions at a fixed friction coefficient? Second, does the same time-rescaling factor apply to different dynamical observables? Third, can discrepancies in the time mapping be reduced or eliminated simply by changing the Langevin friction coefficient, *i.e.,* by moving further into the overdamped regime?

The chain reconfiguration time,*τ_r_*, a fast molecular process directly accessible from single-molecule FRET experiments combined with nanosecond fluorescence correlation spectroscopy (nsFCS)^71,72^, was calculated from the autocorrelation function of the scalar distance between the experimentally labeled residues of ProTα in the CG simulations [Eq. 7-9]. With the resulting value of ∼0.47 ns at 128 mM ionic-strength, the time-rescaling factor, defined as *S_t_* = *τ_exp_*/*τ_CG_*, required to reproduce the experimental reconfiguration time^3,14^ of 14 ns was thus about 30. The ProTα dynamics in the H1–ProTα dimer require rescaling by a factor of 170, and ProTα in the dense phase at 128 mM ionic strength requires rescaling by a factor of 300 [Fig. 4(a)]. The time rescaling factors thus differ by approximately an order of magnitude between isolated ProTα and the dense phase. This result clearly shows that a single global time-rescaling factor is insufficient to map CG simulation dynamics onto experimental dynamics across different molecular environments. The difference in rescaling factors likely arises because both the effective solvent friction and the protein contribution to friction differ between the dilute and dense phases^29,32,73^. In the dense phase, the higher interaction density and multivalent intermolecular contacts modify chain relaxation relative to isolated chains in solution^3,4^. Consequently, applying the same rescaling factor obtained by matching the experimental *τ_r_* of ProTα in the dense phase leads to an overestimation of dilute-phase relaxation times by approximately tenfold for isolated ProTα and approximately twofold for ProTα interacting with H1.

**FIG. 4.**
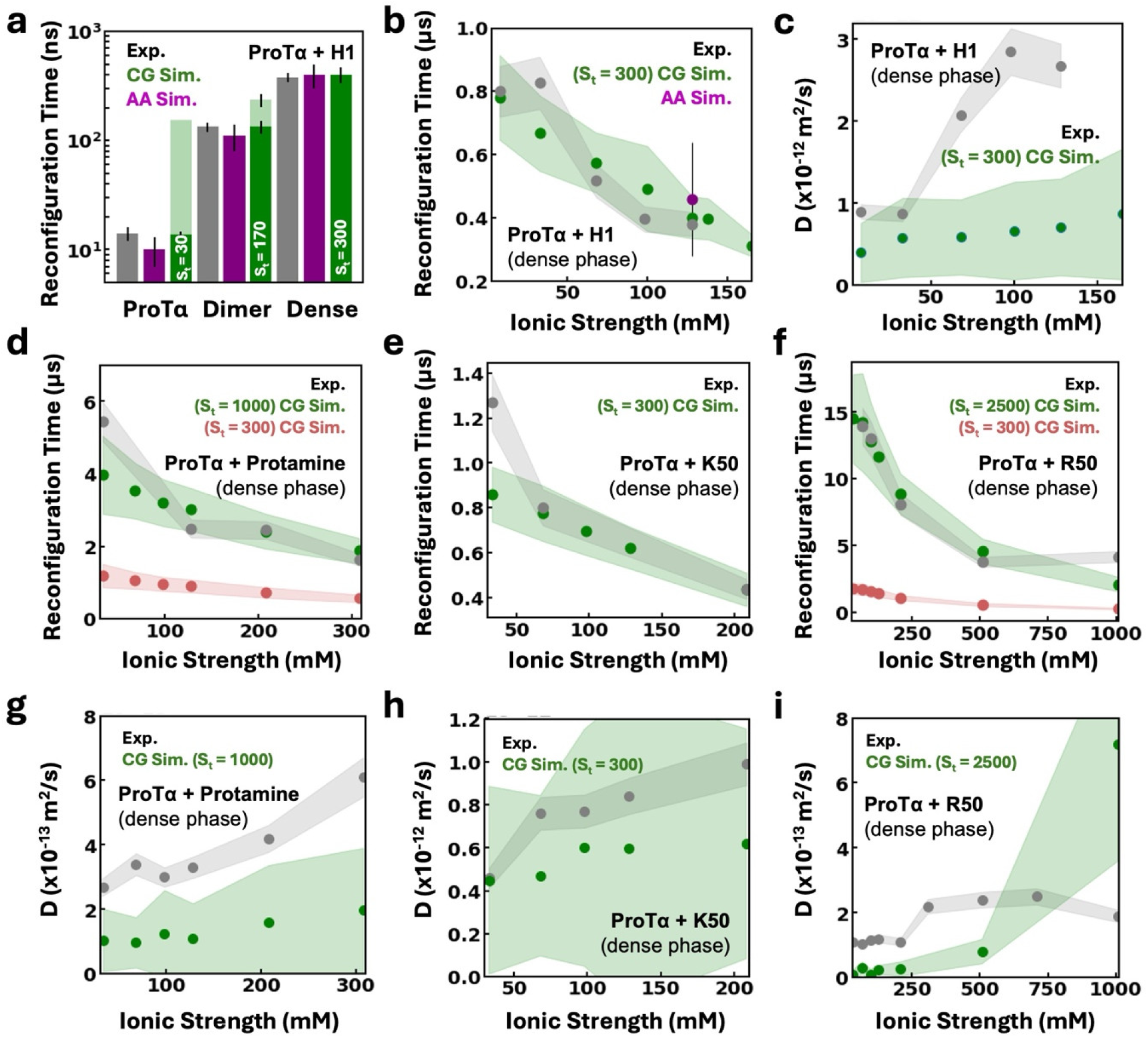
Time rescaling required for molecular dynamics from CG simulations. (a) Comparison of chain reconfiguration time (*τ_r_*) of ProTα in the monomer, dimer, and dense phase with H1. Experimental and AA simulation values are shown as gray and purple bars, respectively. The CG simulation results with a uniform time-rescaling factor (*S_t_*=300) are shown in light green, and the results with a simulation-dependent time-rescaling factor chosen to achieve agreement with experiment are shown in dark green. (b,c) Comparison of *τ_r_* (b) and translational diffusion coefficient *D* (c) of ProTα in complex coacervates with H1, as a function of ionic strength. For the CG simulations, diffusion coefficients were obtained from the center-of-mass MSD in the *y*- and *z*-directions, parallel to the slab interface; motion along the slab-normal *x*-direction was excluded because it is confined by the finite dense-phase thickness. Experimental values^4^ are shown in gray, and CG estimates in green. Shaded bands represent the associated uncertainties, calculated in CG simulations as the standard deviation across different protein chains. (d-f) Same as (b) but for complex coacervates of ProTα with other cationic partners. *τ_r_* calculated using the same rescaling factor as estimated for H1–ProTα condensate is shown in brown. (g-i) Same as panel (c) but for coacervates of ProTα with other cationic partners. All simulations were performed with *γ*=0.2 ps^-1^.

Interestingly, however, within the dense phase, the same time rescaling factor captures the experimentally observed decrease in *τ_r_* with increasing ionic strength^4^ [Fig. 4(b)]. This observation suggests that, over the ionic-strength range examined, the local molecular environment remains sufficiently similar for the same rescaling to apply. The relative dependence of *τ_r_* on ionic strength is consistently reproduced across the four coacervates we simulated [Fig. 4(d)–4(f)]. At higher ionic strengths, chain reconfiguration is faster, consistent with weakened intermolecular electrostatic interactions due to enhanced charge screening. Coacervates formed with lysine-rich cationic partners share the same time-rescaling factor, but reproducing the absolute *τ_r_* of ProTα in coacervates formed with arginine-rich cationic partners requires time rescaling factors that are at least threefold greater. Therefore, the required time rescaling differs not only between the dense and dilute phases but also between systems with different sequence composition, i.e., different interaction parameters between residues [Fig. 4(d) and 4(f)]. Using the reconfiguration-time calibration, the 3 µs CG production simulations correspond to approximately 0.1–7.5 ms of effective physical time, depending on the system, illustrating the much more extensive sampling possible in CG simulations compared to all-atom models.

The translational diffusion coefficient, *D*, of protein chains is another observable available from experiment^4^. Because motion normal to the slab is confined, diffusion coefficients from the CG simulations were calculated from the *y*- and *z*-components of the ProTα center-of-mass, MSD, corresponding to directions parallel to the slab interfaces [Fig. S1]. Using the same system-specific time-rescaling factors as for *τ_r_* would result in the absolute diffusion coefficients being underestimated compared to experiment, but again a single rescaling factor reproduces the relative increase in *D* with increasing ionic strength [Fig. 4(c), 4(g)–4(i)]. Quantitative interpretation of this comparison requires caution. Diffusion coefficients are substantially more difficult to estimate precisely than the intramolecular reconfiguration times considered above. The finite CG trajectories sample heterogeneous local environments, the resulting MSDs can exhibit substantial chain-to-chain variation, and estimates of their long-time slopes are sensitive to the available fitting range. In addition, finite-size effects and the slab geometry can influence translational diffusion^46,74^. We therefore regard the simulated values as effective or apparent lateral diffusion coefficients over the accessible trajectories rather than highly precise estimates of the asymptotic bulk diffusion coefficients. Nevertheless, the rescaled diffusion coefficients remain within the same order of magnitude as the experimental values and qualitatively capture the experimentally observed increase in *D* with increasing ionic strength [Fig 4(c), 4(g)-4(i), and Fig. S2]^4^.

Time-rescaling factors inferred independently from diffusion by optimizing agreement with experiment are approximately 1.5- to 5-fold smaller than those obtained from the reconfiguration times [Fig. S2]. This difference is consistent with the possibility that a single dynamical rescaling factor may not describe all observables equally well^29,32,34,73^. However, given the larger statistical and systematic uncertainties associated with estimating diffusion in the finite-slab simulations, the present data do not allow us to unequivocally attribute the discrepancy to the intrinsic observable dependence of the coarse-grained time mapping^35^.

All comparisons above were performed using the production Langevin friction coefficient of *γ*=0.2 ps^-1^. We therefore tested the extent to which the non-transferability of the dynamical time mapping could be influenced by this particular choice of friction coefficient. We repeated the simulations over a broad range of *γ* values for ProTα as an isolated chain, in the H1–ProTα dimer, and in the H1–ProTα dense phase, as well as for the four dense-phase coacervates. The mean transfer efficiencies remained independent of *γ* in both dilute- and dense-phase systems [Fig. 5(a)-5(b)]. The dense-phase protein concentrations were similarly insensitive to *γ* [Fig. 5(c)], showing that varying the Langevin friction coefficient does not appreciably perturb either the equilibrium chain dimensions or the phase density. This is expected because the Langevin friction controls how rapidly the system explores configuration space, but when consistently coupled to the stochastic force through the fluctuation–dissipation relation, it does not change the canonical equilibrium distribution determined by the conservative interaction potential^75^. Thus, varying the Langevin friction has no observable effect on the equilibrium conformational observables and dense-phase concentrations examined here. Consistency of the simulations with these theoretical expectations provides an additional check on the accuracy of the Langevin integration.

**FIG. 5.**
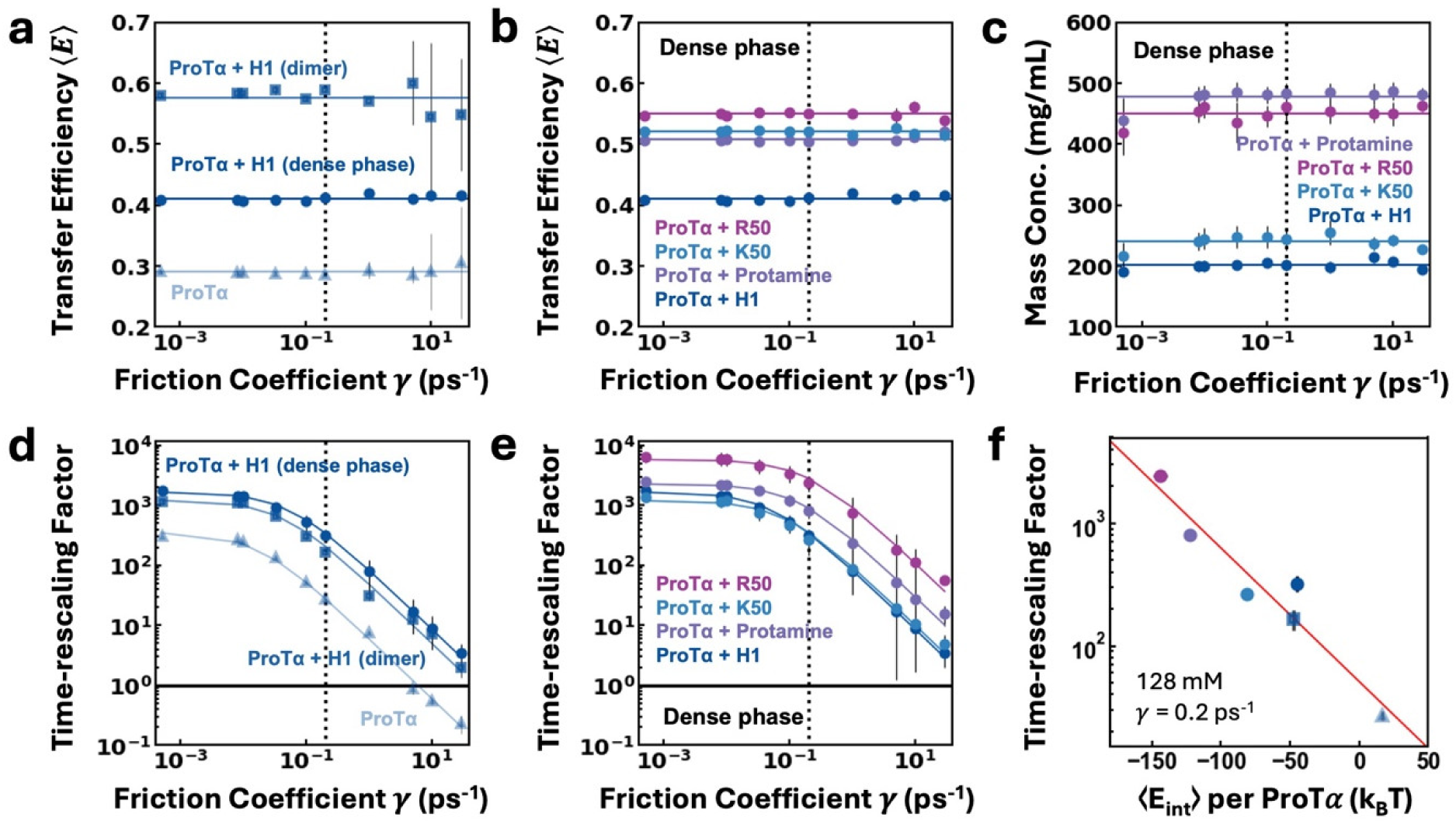
Dependence of equilibrium properties and time rescaling on the Langevin friction coefficient. (a) Mean transfer efficiency (*E*⟩ of ProTα as an isolated chain, in the H1–ProTα dimer, and in the H1–ProTα dense phase as a function of the friction coefficient. (b) Mean transfer efficiency of ProTα in dense phases formed with H1, protamine, K50, and R50. (c) Dense-phase protein concentrations of the four condensates as a function of the friction coefficient. (d) Time-rescaling factors required to reproduce the experimental ProTα reconfiguration times for the three molecular environments shown in (a). (e) Corresponding time-rescaling factors for ProTα in the four dense-phase condensates shown in (b). Solid lines in (d,e) are fits to the empirical shifted inverse function [Eq. (10)]. The dotted vertical line indicates the friction coefficient,*γ*=0.2 ps^-1^, used for the analyses in Fig. 4; the horizontal line at a time-rescaling factor of 1 indicates direct correspondence between CG and experimental time. (f) Time-rescaling factor at *γ* = 0.2 ps̅¹ as a function of the mean total non-bonded interaction energy per ProTα molecule (*E*_int_⟩ at 128 mM ionic strength, for the four condensates, the ProTα–H1 dimer and ProTα monomer (symbols as in panels a-e). More negative (*E*_int_⟩ corresponds to stronger overall favorable non-bonded interactions. The red line is an exponential fit, equivalent to a linear relation between the logarithm of the rescaling factor and (*E*_int_⟩.

However, in contrast to transfer efficiencies and dense-phase concentrations, the time-rescaling factors required to reproduce the experimental reconfiguration times depended strongly and nonlinearly on *γ* [Fig. 5(d)–5(e)]. This dependence was well described by the empirical shifted inverse function [Eq. (10)]. At high friction, *S_t_* approaches the expected *S_t_* ∝ *γ*^−1^ dependence of overdamped Langevin dynamics^50,51^, while at very low friction it deviates from this scaling and exhibits a turnover consistent with the low-friction regime of Kramers dynamics^51^. The results also show that the system-dependent differences in required time rescaling persist over the whole range of friction. Thus, adjusting a single uniform Langevin friction coefficient cannot improve the transferability of the time mapping across systems. Interestingly, the value of 0.2 ps^-1^ used for most of this work appears already close to the overdamped scaling regime where dynamics scales as *γ*^−1^, and so further increasing *γ* changes the overall rate at which each CG system traverses its equilibrium ensemble, but cannot remove the relative dynamical differences between molecular environments or condensate compositions. This observation suggests that the system dependence of time mapping contains a contribution beyond the externally imposed Langevin friction, arising from intermolecular interactions and microscopic degrees of freedom that have been integrated out during coarse-graining.

It might be expected that the system dependence of *S_t_* is related to the strength of molecular interactions of the ProTα chain. We thus compared the time-rescaling factors at the production friction coefficient (*γ*=0.2 ps^-1^) with the mean total non-bonded interaction energy per ProTα chain (*E*_int_⟩ [Fig. 5(f)]. Here, (*E*_int_⟩ includes both intramolecular interaction energy, and intermolecular interaction energy with the surrounding molecules in the dense phase for a ProTα chain, such that increasingly negative values correspond to stronger, more favorable non-bonded interactions. The time-rescaling factor increases as (*E*_int_⟩ becomes more negative. Because the rescaling factor is plotted on a logarithmic scale, the approximately linear dependence in Fig. 5(f) is consistent with an exponential relationship between the required dynamical rescaling and the magnitude of the total non-bonded interactions.

This correlation can be interpreted phenomenologically in a Kramers-like barrier-crossing picture^51^. In Kramers theory^76^, the time of escape from a free-energy minimum contains an activated contribution of the form *τ* ∝ exp (Δ*G*^↕^/*k_B_T*), where Δ*G*^↕^ is the effective free-energy barrier along the relevant reaction coordinate. Coarse-graining removes side-chain orientations, solvent rearrangements, and other microscopic degrees of freedom that can contribute to local barriers for breaking and exchanging intermolecular contacts. If the missing contribution to the effective reconfiguration barrier scales approximately with the overall non-bonded stabilization energy, such that ΔΔ*G*^↕^ ≈ −**Λ**(*E*_int_⟩, then the time-rescaling factor is expected to scale as *S_t_* ∝ exp (−**Λ**(*E*_int_⟩). Here **Λ** is a phenomenological proportionality factor. In this interpretation, stronger non-bonded interactions are associated with a higher microscopic cost of reorganizing the chain and its surrounding contact network at atomistic resolution, whereas the isotropic effective CG interactions smooth part of this microscopic landscape, allowing chains to reconfigure more rapidly. We emphasize that (*E*_int_⟩ is an equilibrium interaction energy and not the activation free energy for chain reconfiguration. The correlation in Fig. 5(f) should therefore not be interpreted as implying that the total non-bonded interaction energy constitutes the barrier to chain reconfiguration. Rather, (*E*_int_⟩ serves as an empirical energetic proxy for the overall strength of interactions, whose microscopic rearrangements may contribute to chain friction.

## **IV.** CONCLUSIONS

We have evaluated the performance of implicit-solvent, residue-level coarse-grained simulations with the HPS model for complex coacervates of charged IDPs. In these models, amino acids are represented as spherical beads, leading to isotropic interactions among neighboring residues in compact IDP ensembles^26,29^. Such interactions are necessarily simplified relative to the chemically specific, anisotropic side-chain interactions present at all-atom resolution. Nevertheless, our simulations reproduce the experimentally observed dense-phase concentrations and their ionic-strength dependence for four complex coacervates of different compositions, while the dilute-phase concentrations are broadly consistent with experiment within their larger statistical uncertainties. Similar agreement in dense-phase concentrations and relative salt-dependent phase stability across the same four condensates has recently been obtained with the independently parameterized Mpipi-Recharged model^13,62^, despite its use of substantially different functional forms for the residue–residue interaction potentials. The consistency between these different coarse-grained models suggests that the experimentally observed phase behavior can be captured by different effective representations of the underlying molecular interactions. Beyond phase diagrams, the present simulations also reproduce the experimentally measured FRET efficiencies reporting on ProTα chain dimensions across dilute and dense phase environments, providing an additional molecular-scale test of the conformational ensembles generated by the coarse-grained model.

The ability of the CG simulations to capture these equilibrium properties suggests that the isotropic interactions between spherically approximated amino acid beads can be interpreted as effective, thermally averaged representations of the underlying anisotropic side-chain interactions. However, equilibrium potentials of mean force in agreement with experimental observables do not necessarily imply that CG simulations reproduce memory/friction effects; dynamic consistency requires additional calibration or the inclusion of generalized Langevin terms^29,32,38^. Experimentally validated all-atom molecular dynamics simulations exhibit longer ProTα residue contact lifetimes in protamine–ProTα coacervates (∼22 ns) than in H1–ProTα coacervates (∼3 ns)^3,4^. This motivates the hypothesis that if the isotropic CG interactions represent an ergodic average over side-chain configurations, then this ensemble-averaged description can also be interpreted as a time-averaged representation of the underlying atomistic interactions. Consistent with this picture, the HPS interaction parameter *λ* is larger for Arg than for Lys^11,16^, reflecting stronger effective interactions and qualitatively paralleling the longer contact lifetimes observed for Arg-mediated interactions in all-atom simulations^4,60^. The time-rescaling factor required to quantitatively reproduce ProTα reconfiguration dynamics may therefore depend on the identity and persistence of these underlying interactions. In line with this expectation, the time-rescaling factor required to quantitatively match the reconfiguration time of ProTα in the dense phase is larger for protamine–ProTα than for H1–ProTα. The same trend is observed for the K50–ProTα and R50–ProTα systems, in which the length and charge patterning of the cationic partners are identical, so the only difference is the identity of the charged residue. This comparison further suggests that residue-specific side-chain chemistry, particularly the distinction between lysine- and arginine-mediated interactions, contributes substantially to the system-specific time-rescaling factors^29,32,73^.

The correlation between the time-rescaling factor and the mean total non-bonded interaction energy in Fig. 5(f) provides an energetic perspective on the same trend. In a Kramers-like interpretation, stronger overall non-bonded stabilization may be associated with larger microscopic barriers or friction associated with rearranging intra- and intermolecular contacts, contributions that are partially smoothed when side-chain and solvent degrees of freedom are integrated out in the CG representation. The resulting relationship should be regarded as phenomenological because (*E*_int_⟩ is an equilibrium interaction energy rather than an activation free energy. Nevertheless, it suggests that the overall non-bonded interaction energy may serve as a useful predictor of the degree of dynamical acceleration introduced by coarse-graining.

Although the Langevin friction coefficient strongly controls the absolute dynamical timescale in the CG simulations, varying γ by several orders of magnitude does not eliminate differences in time rescaling across molecular environments or condensate compositions. At high friction, the systems share the expected *S_t_* ∝ *γ*^−1^dependence, indicating that increasing friction primarily rescales the overall dynamical timescale. However, system-dependent low-friction limits and crossover friction coefficients persist, so adjustment of a single uniform Langevin friction coefficient cannot make time rescaling transferable across systems. These findings highlight that agreement of equilibrium dense-phase concentrations and configurations is not sufficient to infer quantitatively accurate dynamics. Instead, interpreting diffusion, reconfiguration times, viscosity, and molecular exchange in CG condensate simulations requires careful consideration of how coarse-graining modifies the effective dynamical timescale^29,34,73^.

These results connect residue-level CG simulations to polymer-level descriptions of condensate dynamics. In polymer theories, microscopic interactions are often absorbed into effective parameters such as friction, viscosity, or entanglement density ^50,77–79^. Our simulations show how such parameters may emerge from residue chemistry: lysine- and arginine-rich condensates can exhibit similar equilibrium chain dimensions while requiring different time-rescaling factors to reproduce molecular dynamics. This observation suggests a possible link between interaction energies and the effective internal friction, or to the renormalized segmental mobility, in polymer models of coacervates^80,81^. Similarly, effective diffusion or friction along the folding coordinates can encode microscopic transitions and landscape roughness, providing a precedent for interpreting time-rescaling factors as emergent dynamical parameters^35,51^.

More broadly, the scope of this manuscript extends beyond charged biomolecular condensates by placing residue-level CG dynamics in the same conceptual framework used for folded proteins and soft-matter liquids. In folded proteins, CG models can often preserve native-state structure, thermodynamic stability, and conformational preferences while altering barrier-crossing kinetics, side-chain packing relaxation, and internal friction, because solvent and fast side-chain degrees of freedom have been averaged out^26,27,30,43,82–84^. Recent work on protein folding and unfolded-chain dynamics further shows that internal and memory-dependent friction can play a central role in determining molecular kinetics, underscoring that the free-energy landscape alone is insufficient to specify dynamical timescales^40,42,43,85,86^. In soft-matter liquids and polymer melts, analssogous coarse-graining commonly accelerates dynamics and can require state-dependent, and in some cases observable-dependent, time mapping, memory kernels, or friction renormalization to recover transport and rheological properties^29,31,32,34,77,87^. The present condensate simulations connect these two limits, showing that residue-specific interaction potentials in CG models can preserve equilibrium observables, such as phase density and chain dimensions, while yielding different dynamical timescales because coarse-graining integrates out microscopic contact dynamics, local friction, and memory effects. These inferences show that dynamically consistent CG models of biomolecular condensates should not rely on a single global time-rescaling factor. A useful next step would be to combine thermodynamic validation with kinetic calibration against reconfiguration, diffusion, exchange rates, and viscosity. Predictive models may ultimately need to encode sequence- and chemistry-dependent friction, hydrodynamic screening, or generalized Langevin memory terms^4,29,38,42,88^, which might be related to interaction energies. Such connections might help convert empirical time-rescaling factors into predictive, sequence-dependent friction parameters for biomolecular condensates^31,32,34,87,89^.

## SUPPLEMENTARY MATERIAL

The supplementary material contains amino acid sequences of the simulated polypeptides (Table S1) and additional analyses of translational diffusion coefficients (Figs. S1 and S2).

## ACKNOWLEDGMENTS

We thank Miloš T. Ivanović for discussions and assistance with some of the CG simulations. This work was supported by the Swiss National Science Foundation (to B.S., 310030_197776 and CRSII5_205922), and the Forschungskredit of the University of Zurich (00109396 to S.G.). R.B.B. was supported by the Intramural Research Program of the National Institute of Diabetes and Digestive and Kidney Diseases (NIDDK) within the National Institutes of Health (NIH). The contributions of the NIH authors are considered Works of the United States Government. The findings and conclusions presented in this paper are those of the authors and do not necessarily reflect the views of the NIH or the U.S. Department of Health and Human Services. We used the computational resources of Piz Daint, Alps, and Eiger at the Swiss National Supercomputing Centre (CSCS), and of the National Institutes of Health HPC Biowulf cluster (http://hpc.nih.gov).

## AUTHOR DECLARATIONS

### Author Contributions

**Soundhararajan Gopi:** Conceptualization, Methodology, Software, Validation, Formal analysis, Investigation, Visualization, Supervision, Writing – original draft, Writing – review & editing. **Hanling Qin:** Investigation, Formal analysis, Visualization, Writing – review & editing. **Robert B. Best:** Conceptualization, Methodology, Software, Supervision, Writing – review & editing. **Benjamin Schuler:** Conceptualization, Supervision, Funding acquisition, Project administration, Writing – review & editing.

## DATA AVAILABILITY

The data that support the findings of this study are available from the corresponding author upon reasonable request.

