## Supplementary File for "Beyond Equilibrium Ensembles: Time Rescaling in Coarse-Grained Simulations across Single-Molecule and Condensate Regimes"

| | |
|---|---|
| ProT$\alpha$ | PSDAAVDTSSEITTKDLKEKKEVVEEAENGRDAPANGNAENEENGEQEADNEVDEEEEEGGEEEEEEEEGDGEEEDGDEDEEAESATGKRAAEDDEDDDVDTKKQKTDEDD |
| H1 | CTENSTSAPAAKPKRAKASKKSTDHPKYSDMIVAAIQAEKNRAGSSRQSIQKYIKSHYKVGENADSQIKLSIKRLVTTGVLKQTKGVGASGSFRLAKSDEPKKSVAFKKTKKEIKKVATPKKASKPKKAASKAPTKKPKATPVKKAKKKLAATPKKAKKPKTVKAKPVKASKPKKAKPVKPKAKSSAKRAGKKKGGPR |
| Protamine | MPRRRRSSSRPVRRRRRPRVSRRRRRRGGRRRR |
| Poly-L-lysine 50 (K50) | KKKKKKKKKKKKKKKKKKKKKKKKKKKKKKKKKKKKKKKKKKKKKKKKKK |
| Poly-L-arginine 50 (R50) | RRRRRRRRRRRRRRRRRRRRRRRRRRRRRRRRRRRRRRRRRRRRRRRRRR |

**Table S1.** Amino acid sequences of polypeptides used in simulations.

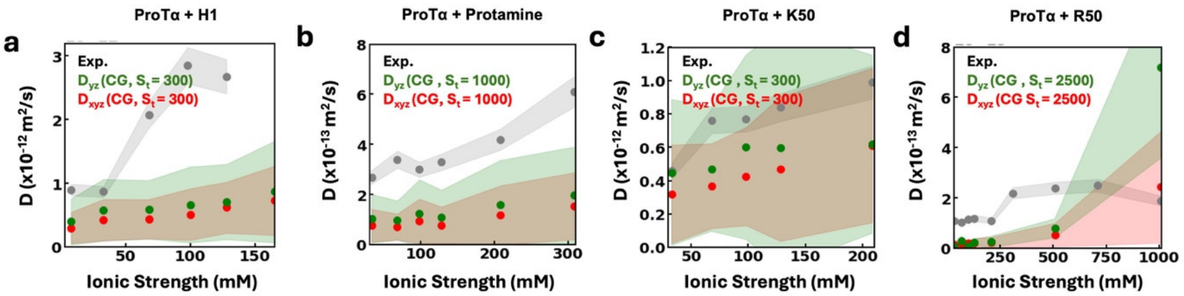


**FIG. S1.** Diffusion coefficients $D$ are shown as a function of ionic strength for complex coacervates of (a) ProTα + H1, (b) ProTα + protamine, (c) ProTα + K50, and (d) ProTα + R50. Experimental diffusion coefficients[1] are shown in gray, while coarse-grained (CG) estimates obtained from the full three-dimensional displacement ($D_{xyz}$) are shown in red. Green symbols indicate the in-plane diffusion coefficient calculated ($D_{yz}$) from motion in the *yz* plane. Shaded regions indicate the corresponding uncertainty ranges. The time-rescaling factor, $S_t = \tau_{\text{exp}}/\tau_{\text{CG}}$, is specified in the figure legends.

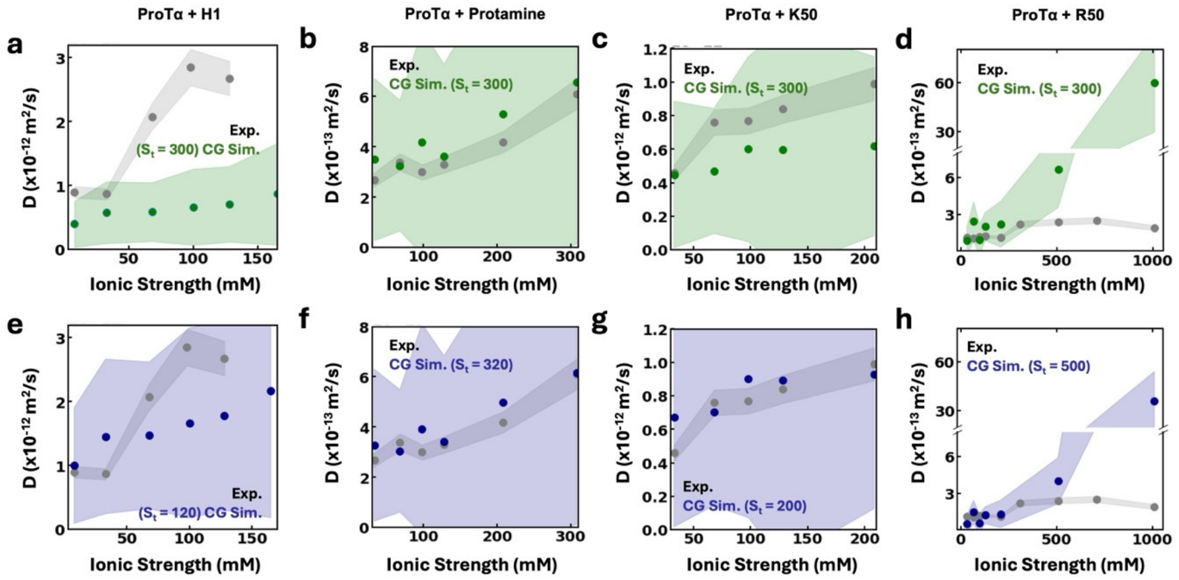


**FIG. S2.** Comparison of diffusion coefficient $D$ as a function of ionic strength for the same systems as in Fig. S1 estimated from the CG simulations (green), along with the data from single-molecule experiments[1] (gray), assuming the same time rescaling factor for all simulations (a-d) and using a system-dependent time rescaling factor (blue) to maximize the agreement with experimental data (e-h). The time-rescaling factor, $S_t = \tau_{\text{exp}}/\tau_{\text{CG}}$, is specified in the figure legends.
